# GOLEM: A tool for visualizing the distribution of Gene regulatOry eLEMents within the plant promoters with a focus on male gametophyte

**DOI:** 10.1101/2024.08.05.606583

**Authors:** Lukáš Nevosád, Božena Klodová, Tomáš Raček, Tereza Přerovská, Alžbeta Kusová, Radka Svobodová, David Honys, Petra Procházková Schrumpfová

## Abstract

**Background:** The regulation of gene expression during tissue development is very complex. A key mechanism of gene regulation is the recognition of regulatory motifs, also known as cis-regulatory elements (CREs), by various proteins in gene promoter regions. Localization of these motifs near the transcription start site (TSS) or translation start site (ATG) is crucial for transcription initiation and rate. Transcription levels of individual genes, regulated by these motifs, can vary significantly across tissues and developmental stages, especially in processes like sexual reproduction. However, the precise localization and visualization of regulatory motifs in relation to gene expression in specific tissues can be challenging.

**Results:** Here, we introduce a program called GOLEM (Gene regulatOry eLEMents) which enables users to precisely locate any motif of interest with respect to TSS or ATG within the relevant plant genomes across the plant Tree of Life (*Marchantia, Physcomitrium, Amborella, Oryza, Zea, Solanum* and *Arabidopsis*). The visualization of the motifs is performed with respect to the transcript levels of particular genes in leaves and male reproductive tissues and can be compared with genome-wide distribution regardless of the transcription level. Additionally, genes with specific CREs at defined positions and high expression in selected tissues can be exported for further analysis. GOLEM’s functionality is illustrated by its application to conserved motifs (e.g. TATA-box, ABRE, I-box, and TC-element), as well as to male gametophyte-related motifs (e.g. LAT52, MEF2, ARR10_core, and DOF_core).

**Conclusion:** GOLEM is a freely available tool (https://golem.ncbr.muni.cz) for tracking the precise localization and distribution of any CREs of interest in plant gene promoters.

**Graphical abstract:** 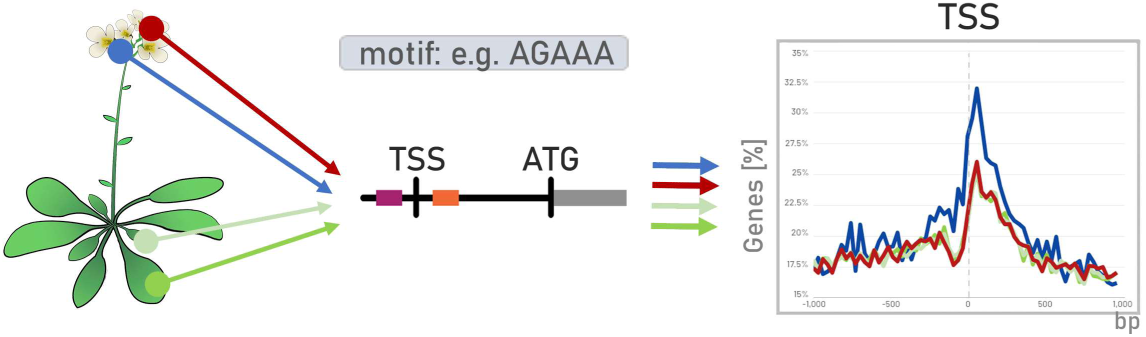

## 1 Introduction

The regulation of gene expression is a dynamic process that requires a tightly orchestrated control mechanism. This regulation is crucial not only for maintaining proper cellular function but also for precise regulation of gene transcription and control of cell differentiation into specific tissues and organs. Cis-regulatory elements (CREs) are short DNA sequence motifs that act as molecular switches activating or repressing gene expression [1,2]. In order to do that, CREs serve as binding sites for various regulatory proteins, including transcription factors (TFs) [3].

When performing CREs localization within the plant or animal genomes, FIMO (Find Individual Motif Occurrences) or CentriMoLocal (Motif Enrichment Analysis) that are part of the MEME-suit [4,5] are popular tools. However, users must consider certain limitations, such as uploading input already pre-processed data in specific formats and understanding proper parameter settings. Moreover, analyzing DNA sequences from promoter regions relevant to genes with high transcription in specific tissues, based on RNA-seq data, can be challenging for those lacking bioinformatics expertise, even when using the web interface with preconfigured settings. Additionally, inconsistencies in gene nomenclature systems in many plant species [6–9], compared to well-established model organisms (e.g., thale cress, human), can pose obstacles for automatization of these procedures.

Transcriptomics has gained significant popularity in recent years, as it can provide detailed insights into gene expression dynamics across different tissues, developmental stages, or experimental conditions [10]. Transcriptome sequencing (RNA-Seq) is an important method for investigating gene regulation. While most transcriptome analyses have traditionally focused on easily accessible plant materials such as leaves or seedlings, there is a growing trend toward exploring transcriptomes from intricate and deeply embedded tissues, including sperm cells and various pollen developmental stages [11,12]. These innovations, together with advances in bioinformatics, are facilitating breakthroughs in understanding the regulation of plant reproduction. By determining overrepresented CREs in the promoters of differentially expressed genes, the candidate tissue-specific transcriptional regulators can be identified [13].

The first identified eukaryotic CRE within the gene promoters, owing to its predictable locations surrounding gene transcription start sites (TSS), was TATA-box (TATAWA) [14–16]. The TATA-box has been conserved throughout the evolution of eukaryotes and is usually located 25-35 base pairs (bp) upstream of TSS [17]. Even though the TATA-box is a common CRE, it is not a general feature of all promoters. Only a small fraction of eukaryotic genes actually harbor a TATA-box: 20% to 46% of promoters in yeast [18,19] or less than 10% of genes in human [20,21]. In plants, less than 39% of thale cress (*Arabidopsis thaliana*) promoters contain a TATA-box or a TATA-variants [22,23], whereas approximately 19% of rice (*Oryza sativa*) genes possess the TATA-box [24]. In recent years, it became apparent that there are no universal promoter elements across species and some promoter elements are involved in enhancer promoter specificity as well as specific biological networks. However, precise regulatory sequences at precise locations are essential for promoter function [25,26].

Specific CREs can activate the expression of particular target genes [26]. For instance, the core sequence GATY is recognized by type-B Arabidopsis Response Regulators (ARR10-B), proteins that mediate the cytokinin primary response [27,28]. The I-box is involved in light-regulated and/or leaf-specific gene expression of photosynthetic genes [29–31]. The I-box motifs (GATAAG) were found in most of the genes crucial for plant photosynthesis [29,30].

Plant sexual reproduction possesses an extraordinary ability to establish new cell fates throughout their life cycle, in contrast to most animals that define all cell lineages during embryogenesis [32]. Sexual reproduction was introduced after the origin of meiosis [33] and the life cycle, in which diploid sporophyte alternate with the haploid gametophyte in land plants [34]. The key mediators of developmental and organismal phenotypes are CREs, that can orchestrate precise timing and magnitude of gene transcription, especially during the development of male or female gametophyte tissues [35]. Several CREs involved in the regulation of key genes required for the differentiation of male germline, active in sperm cells and pollen vegetative cells, have already been identified: MEF2-type CArG-box (CTA(A/T)_4_TAG [36]); LAT52 pollen-specific motif in tomato (AGAAA [37]); DOF core motif (AAAG [38,39]); and many others [38,40–42]. However, the precise distribution of these motifs within promoters, particularly their proximity to TSS or translation start site (ATG), and their prevalence in the promoters of genes exhibiting higher transcription levels in specific tissues related to plant reproduction, remain unclear.

Here we present a user-friendly online software GOLEM (Gene regulatOry eLEMents) https://golem.ncbr.muni.cz, which allows browsing various tissues such as sporophyte (leaves) or male gametophyte developmental tissues (antheridia, pollen stages, sperm cells) across the selected plant genomes within the plant Tree of Life (mosses, basal angiosperm, monocots and dicots). Our software enables us to investigate the precise localization and distribution of any CREs of interest in gene promoters, in proximity to the TSS and ATG. The set of investigated genes can be specified by the level of gene expression in specific tissues based on transcriptomic data. Furthermore, tracking of the genome-wide distribution across exemplified genomes, regardless of the transcription level, may aid to track the evolution of regulatory motifs across the plant Tree of Life. Finally, a set of genes with only specific CREs at defined positions showing high expression only in the tissue of interest can be exported for further analysis, including for instance protein functional enrichment analysis. We demonstrate the utilization of the GOLEM program not only on motifs associated with male gametophyte development, such as LAT52, MEF2, ARR10_core, and DOF_core, but also on conserved motifs such as the TATA-box, ABRE, TC-element, I-box and DRE/CRT element.

## 2 Materials and Methods

The GOLEM program is divided into two main phases: data processing pipeline and data visualization.

### Data processing pipeline

#### Segmentation of genomic sequences upstream/downstream of TSS and ATG

The reference genomes and genome annotations files from *Marchantia polymorpha*, *Physcomitrium patens*, *Amborella trichopoda*, *Oryza sativa*, *Zea mays*, *Solanum lycopersicum*, and *Arabidopsis thaliana* were downloaded in the FASTA format and General Feature Format (GFF3), respectively (**Supplementary Table 1A**). The data processing pipeline first parses location data of individual genes on a reference genome, using annotation data from a GFF3 file, to identify the position of TSS (transcription start site) and ATG (first translated codon) in the reference genome. The locations of TSS and ATG were determined as positions of “five_prime_UTR” and “start codon” in GFF3, respectively. The analyzed dataset comprises a defined segment of genomic sequences specified by the user (e.g., <−1000, 1000> bp) upstream and downstream of the TSS or ATG.

#### TPM values from various plant tissues and developmental stages

The pipeline matches individual genes against a Transcript Per Million (TPM) table from various tissues and developmental stages. The TPM values express normalized transcription rates of individual genes obtained from RNA-seq datasets. The TPM for *Arabidopsis* leaves, seedling, egg, sperm, semi-in vivo grown pollen tube (SIV_PT; hereafter referred to as pollen tube (PT)), and tapetum samples together with all *Zea mays* samples were processed in the following manner: the RNA-seq datasets were downloaded as fastq files from Sequence Read Archive (SRA, https://www.ncbi.nlm.nih.gov/sra/), accession codes are stated in **Supplementary Table 1B**. The raw reads were checked for read quality control (Phred score cutoff 20) and trimmed of adapters with Trim Galore! (https://www.bioinformatics.babraham.ac.uk/projects/trim_galore/, v 0.5.0). Next, the reads were mapped with the use of Spliced Transcripts Alignment to a Reference (STAR) v 2.7.10a [43] aligner to both genome and transcriptome. The TPM values were calculated from RNA-Seq using Expectation Maximization (RSEM) [44] on gene level (*Arabidopsis, Zea).* The TPM values of sample replicas were then averaged and used as an input data for GOLEM. TPM values for *A. thaliana* early pollen stage were calculated as a mean of Uninucleate microspore (UNM) and Bicellular pollen (BCP) stages; for the late pollen stage as a mean of Tricellular pollen (TCP) and Mature pollen grain (MPG) stages. *Arabidopsis* genes were annotated by The Arabidopsis Information Resource (TAIR, https://www.arabidopsis.org/tools/bulk/genes/index.jsp <u>on</u> arabidopsis.org, 10.10.2022), while MaizeMine v.1.5 (https://maizemine.rnet.missouri.edu/maizemine/begin.do) was used for gene annotation of *Zea*. The GFF3 files of all organisms were processed with AGAT analysis toolkit (https://zenodo.org/record/7255559#.ZAn_kS2ZPfY, v1.0.0) prior their usage in GOLEM. The TPM values for *M. polymorpha*, *P. patens*, *A. trichopoda*, *O. sativa*, and *S. lycopersicum* tissue were acquired from Conekt database (https://conekt.sbs.ntu.edu.sg, [11]). The TPM values for pollen developmental stages of *A. thaliana* Columbia-0 (Col-0) and Landsberg erecta (Ler) were extracted from Klodová et al. [12]. The normalized TPM values used for the data processing pipeline are listed in **Supplementary Table 2**.

#### Individual gene matching against TPMs

The pipeline matches individual genes against a TPM table from various tissues and developmental stages (hereafter referred to as stages). The sequential values of a series of TPM numbers are pooled. The output of this data processing pipeline is a separate FASTA-compatible file that contains each valid gene from the original input, along with information about the position of TSS and ATG, and transcription rates (TPMs) in each stage added as comments. The pipeline also generates a validation log that provides information about genes that were excluded, i.e., non-protein coding genes (noStartCodonFound), pseudogenes (noFivePrimeUtrFound, noTpmDataFound), genes without TSS (noFivePrimeUtrFound, if relevant for certain organism). In *A. trichopoda* and *O. sativa*, the GFF3 gene annotation of TSS is inadequate, limiting the search to motifs in the vicinity of the ATG.

#### Motif search

Motif search uses regular expressions to search the input string of base pairs. For each motif, the reverse complement is calculated and then translated together with the forward strand into regular expressions. When the regular expression is run against the input data, we record all results and calculate their relative positions (adjusted to the middle of the motif) relative to TSS and ATG. The motif sequences are searched in the buckets that can be specified by the user (default size is 30 bp).

### Data visualization

The data visualization phase consists of five steps, as depicted in **Fig. 1** and **Supplementary Fig. 1** In the selected genome (**Fig. 1A**) the user can choose the genomic interval to be searched, effectively specifying the window of the sequence where the search is performed, and focus on a defined region in the vicinity of the TSS or ATG (**Fig. 1B** and **Supplementary Fig. 1B**). A single custom motif or multiple motifs, including degenerate motifs of interest can be defined by users (**Fig. 1C** and **Supplementary Fig. 1C**). Optionally the motif can be chosen from several motifs present in the software: i) conserved eukaryotic promoter motif: TATA-box (TATAWA; [14–16]); ii) motifs associated with pollen development: pollen Q-element (AGGTCA; [45]); POLLEN1_LeLAT52 (AGAAA; [37,46]); CAAT-box (CCAATT; [47]); GTGA motif (GTGA; [48]); iii) motifs associated with plant hormone-mediated regulation: ABRE motif (ACGTG; [49,50]); ARR10_core (GATY; [27,51]); E-box (CANNTG; reviewed in [52]); G-box (CACGTG; [53,54]); GCC-box (GCCGCC; [55,56]); iv) biotic and abiotic stress responses: BR_response element (CGTGYG; [57–59]); DOF core motif (AAAG; [38,39]; reviewed in [60]); DRE/CRT element (CCGAC; [61–63]); and v) motifs regulated by light: I-box (GATAAG; [29,30]).

**Fig. 1.**
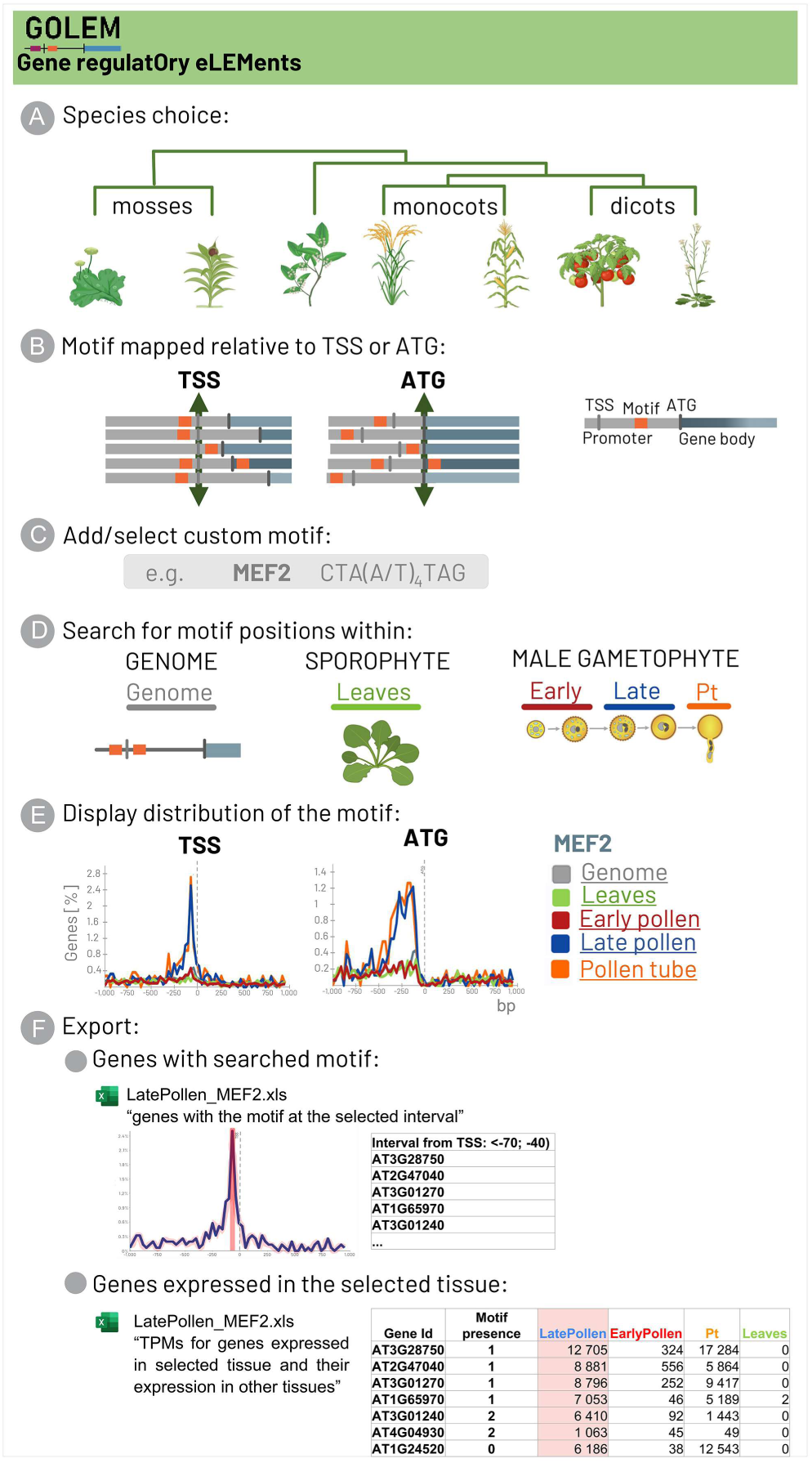
An illustrative overview of the workflow of the GOLEM software. (A) The plant species across the plant Tree of Life is chosen. (B) Region in the vicinity of the transcription start site (TSS) or translation start site (ATG) is specified. (C) Motifs of interest are defined. (D) The promoters of the genes showing expression in selected tissue (sporophyte, male gametophyte), together with an analysis of genome-wide distribution regardless of transcription, are chosen for the analysis. (E) The exemplified MEF2-type CArG-box motif shows distribution upstream of TSS, with higher prevalence in the promoters of genes transcribed during late pollen development. (F) The accession numbers (Gene ID) of genes with certain motif within a defined region and tissue, or genes expressed in selected tissues, may be exported in XLSX format tables.

Motifs are searched in both forward and reverse forms, and the reverse form is calculated automatically (**Supplementary Fig. 1D**). Before the entire analysis, the user confirms the stages to be searched (**Fig. 1D** and **Supplementary Fig. 1E**) and selects the method for choosing genes for analysis (**Supplementary Fig. 1F**). Gene selection, based on TPM, uses a given percentile (genes whose transcripts will represent e. g. 90% of all transcripts transcribed from the total number of protein-coding genes in each selected stage; default is 90^th^ percentile) to select the genes that are the most/least transcribed in tissues or developmental stages of interest, to exclude the genes with low or even negligible transcription. The number of selected genes within the given percentile can be tracked during the proceeding steps. In addition to selecting genes based on a specified percentile of transcription levels, users can choose a specific number of genes with the highest or lowest transcription levels for analysis. The motif distribution, regardless of the transcription level, can be also included into the analysis (stage genome, hereafter referred as “all”).

The goal of the analysis is to visualize the distribution of motifs of interest in the vicinity of the TSS or ATG of all protein-coding genes, or exclusively in selected genes that exhibit high/low transcription levels in particular tissues and developmental stages (**Fig. 1E** and **Supplementary Fig. 1G**). Results are presented graphically, with each stage color-coded, and can be displayed as percentages of the genes with a certain motif or as simple counts. The user can also choose to display either the number of motifs found or aggregate the motifs by genes (**Supplementary Fig. 1H**).

Additionally, the set of genes with specific motifs at defined positions can be exported for further analysis (**Fig. 1F** and **Supplementary Fig. 1I, J**). Within the analysis, the application allows the user to export each data series or export the aggregated data for all data series in XLSX format. The user can also see the distribution of individual motifs and drill down through them.

The data visualization phase involves an application written in Flutter/Dart [64] which can be run as a standalone application or compiled into JavaScript and hosted on the web as a single-page web app (https://golem.ncbr.muni.cz).

### Limitations

The entire processing takes place on the client within the application or web browser, with input files loaded into memory. The program’s ability to work with large datasets may be constrained by the available memory and the web browser’s local client settings. Nevertheless, we found that even in the web application, where performance is limited due to the inefficiencies of JavaScript compared to platform native code, performance is satisfactory on modern computers without the need for significant code optimization.

### Gene ontology annotation and functional analysis

To interpret Gene Ontology (GO), genes containing the motif of interest (LAT52) located between −70 to −10 bp from the ATG start codon and expressed in the 80^th^ percentile during the late pollen stage in *A. thaliana* were exported from the GOLEM in XLSX table. These genes were identified by GOLEM within the interval <−1000, 1000> bp relative to the ATG, using a 30 bp bucket size. The list of AGI locus codes (gene identifiers) was exported and then uploaded to the g:PROFILER software [65,66] for functional annotation analysis, using default parameters. The functional annotation covered biological processes (BP), cellular compartments, and molecular functions. Further, the genes listed under BP in g:PROFILER were uploaded to Search Tool for the Retrieval of Interacting Genes/Proteins (STRING, https://string-db.org, [67,68]), where genes associated with the same biological processes were clustered and highlighted.

## 3 Results and Discussion

### Gene expression dynamics during the male gametophyte development

User-friendly online software GOLEM allows browsing various tissues such as leaves, leaflets, or tissues associated with plant sexual reproduction (antheridia, pollen stages, sperm cells) across the selected plant genomes and investigates the precise localization and distribution of any CREs of interest in gene promoters, in proximity to the TSS and ATG. The set of investigated genes, in each tissue or in individual pollen developmental stages, can be specified by the level of gene expression in specific tissues based on transcriptomic data and calculated values of TPM. However, plant sexual reproduction is a complex process involving specialized structures at several stages that can significantly differ in the level of their transcription [69,70].

In flowering plants during the early stages of male gametophyte (pollen) development, the haploid uninucleate microspore (UNM) divides asymmetrically to form bicellular pollen (BCP), which is comprised of a large vegetative cell and small generative cell in a unique ‘‘cell-within-a-cell’’ structure. In approximately 30% of angiosperms, including *A. thaliana, O. sativa*, *Z. mays* the generative cell divides again to form tricellular pollen (TCP; reviewed in [71]) so that the mature pollen grain (MPG) is tricellular, composed of the vegetative cell and two sperm cells. After reaching the stigma, the growing pollen tube (PT) is guided to the female gametophyte (ovules) to deliver the sperm cells. In 70% of species, including *S. lycopersicum* and basal angiosperm as *A. trichopoda* [72], the MPG is bicellular and becomes tricellular after the MPG reaches the papillary cells of the stigma, where it is rehydrated and activated (reviewed in [70,73]). In bryophytes, such as the liverwort (*M. polymorpha*) and mosses (*P. patens*), the haploid gametophyte generation is the dominant phase of the life cycle. In bryophytes, the male gametophyte is called the antheridia and holds the male gametes (sperm cells; [9,74]).

GOLEM is based on comparing the expression of individual genes across various developmental stages and tissue samples based on TPM. In many angiosperms, including *A. thaliana* or *N. tabacum*, a substantial reduction in the number of expressed genes and significant changes during the transition from early pollen stages (UNM, BCP) to late pollen stages (TCP, MPG) were reported [12,75]. Due to the significant differences in gene expression between the stages, TPMs between various stages cannot be compared directly [76]. To overcome this, the sequential values of a series of TPM numbers are pooled and compared, as this enables comparison across multiple samples with varying numbers of input values. The genes with the highest transcription in each stage can be set as a percentile (e.g., 90^th^ percentile comprises the genes whose transcripts represent 90% of all transcripts transcribed from the total number of protein-coding genes) or as certain number of the genes. The number of the genes in chosen percentile, from total number of the validated genes included into analysis, can be tracked in the GOLEM outputs (**Supplementary Fig. 1G, J**). The results can be further displayed as percentages of the genes with certain motif, however the exact counts of the motifs can be tracked alongside (**Supplementary Fig. 1G, H**).

Our analysis confirmed the reduction in the number of expressed genes at late pollen developmental stages described previously in [12,77,78]. In early pollen stages, leaves and seedlings of *A. thaliana* the genes composing the 90^th^ percentile represent 27%, 24% and 30% of the total protein-coding genes, respectively. In late pollen stages, sperm cells and PT, those genes represent 7%, 3% and 5%, respectively (**Supplementary Fig. 2A**). On the other hand, in bryophyte *M. polymorpha*, the genes whose transcripts comprise the 90^th^ percentile show more similar levels in antheridia, sperm cells and thallus, 25%, 30% and 29%, respectively (**Supplementary Fig. 2B**).

### Positional distribution of peaks reveals preferential localization of the searched motifs to the TSS or ATG

The GOLEM software aligns all genes relative to the TSS or ATG and conducts a comprehensive analysis of the CREs in their proximal regions (**Fig. 1B**). When comparing the results from TSS and ATG-based analyses, it is crucial to consider the variations in the length of the 5’ UTR. The median length of the 5’ untranslated regions (5’UTRs), i.e., the region between the TSS and ATG, is not uniform across the plant species. The median length of 5’UTR is 454 bp in *M. polymorpha* [79]; 477 bp in *P. patens* [80]; 111 bp in *O. sativa* [80,81]; 179 bp in *Z. mays* [80]; 214 bp in *S. lycopersicum* [80]; and 184 bp in *A. thaliana* [80]. However, genes with very short (1-50 bp) or very long (>2000 bp) 5’UTRs were also detected [12,75]. Due to the varying lengths of the 5’UTR, it is possible to determine the positional distribution of the peak near the TSS or ATG (whether it is sharp, narrow, or bell-shaped) [82]. It enables to determine whether the motif of interest is preferentially localized near the TSS or ATG.

The positional distribution and revelation of the preferential motif localization can be exemplified by the TC-elements (TC_(n)_, TTC_(n)_). TC-element is a motif described in *A. thaliana* and *O. sativa* promoters but not in *Homo sapiens* or *Mus musculus*. TC-element is preferentially present in the promoters of genes involved in protein metabolism [22]. The authors showed the peak is centered −33 to +29 bp to TSS. Our detailed analysis in GOLEM showed a peak of TC-element centered −20 bp from ATG (**Supplementary Fig. 1C, G** **and** **Supplementary Fig. 3**), rather than −30 bp upstream TSS as was reported in [22]. Moreover, dehydration-responsive element/C-repeat (DRE/CRT, CCGAC) element [62,83], a CRE detected in promoter regions of several target stress-responsive genes [61,63,84–87], exhibits a bell-shaped peak downstream of the TSS, but a sharp peak downstream of the ATG using GOLEM. This pattern suggests that the DRE/CRT element is preferentially located in the gene bodies, downstream of ATG, rather than in promoters of genes that exhibit higher expression under non-stressed conditions in *A. thaliana*. Similarly, the ABA-responsive cis-element-coupling element1 (ABRE motif, ACGTG), which is involved in the abscisic acid (ABA) responsiveness [49], shows a sharp peak upstream of the TSS, but more bell-shaped peak upstream the ATG in. *A. thaliana* (**Supplementary Fig. 3**), as was previously suggested in *O. sativa* aleurone cells [50].

### GOLEM reveals that TATA-box-containing promoters are associated with late pollen development

Gene expression is primarily controlled through the specific binding of various proteins to diverse DNA sequence motifs upstream/downstream of the TSS [88]. TATA-box is a particularly well-conserved preferentially located motif since it is found in the same promoter region in both plants and animals. In *A. thaliana* and *O. sativa* genomes, the TATA-box is strictly located within the −39, −26 region upstream of the TSS [22]. Even though the TATA-box is common CRE, it is not a general feature of all promoters [18,19,22–24]. Moreover, the percentage of the genes with TATA-box may be associated with level of the expression. In barley embryo, the TATA-box containing promoters are associated mostly with genes exhibiting high expression levels, while promoters lacking a distinct TATA-box tend to exhibit lower expression. The genes regulated by the TATA-box promoters were annotated as responsive to environmental stimuli, stress, and signals related to hormonal, developmental, and organ growth process levels [25].

Our GOLEM program allowed us easily to verify that only a small fraction of plant genes actually harbor a TATA-box (TATAWA) in their promoter in −30 to −25 area upstream of TSS, regardless of the transcription of those genes (**Fig. 2A**), as seen in the plant species with annotated TSS positions. Even though in bryophytes, such as the liverwort (*M. polymorpha*) and mosses (*P. patens*) the percentage of the genes with the TATA-box upstream of TSS is negligible, those genes also show preferential location in −30 to −25 area upstream of TSS.

**Fig. 2.**
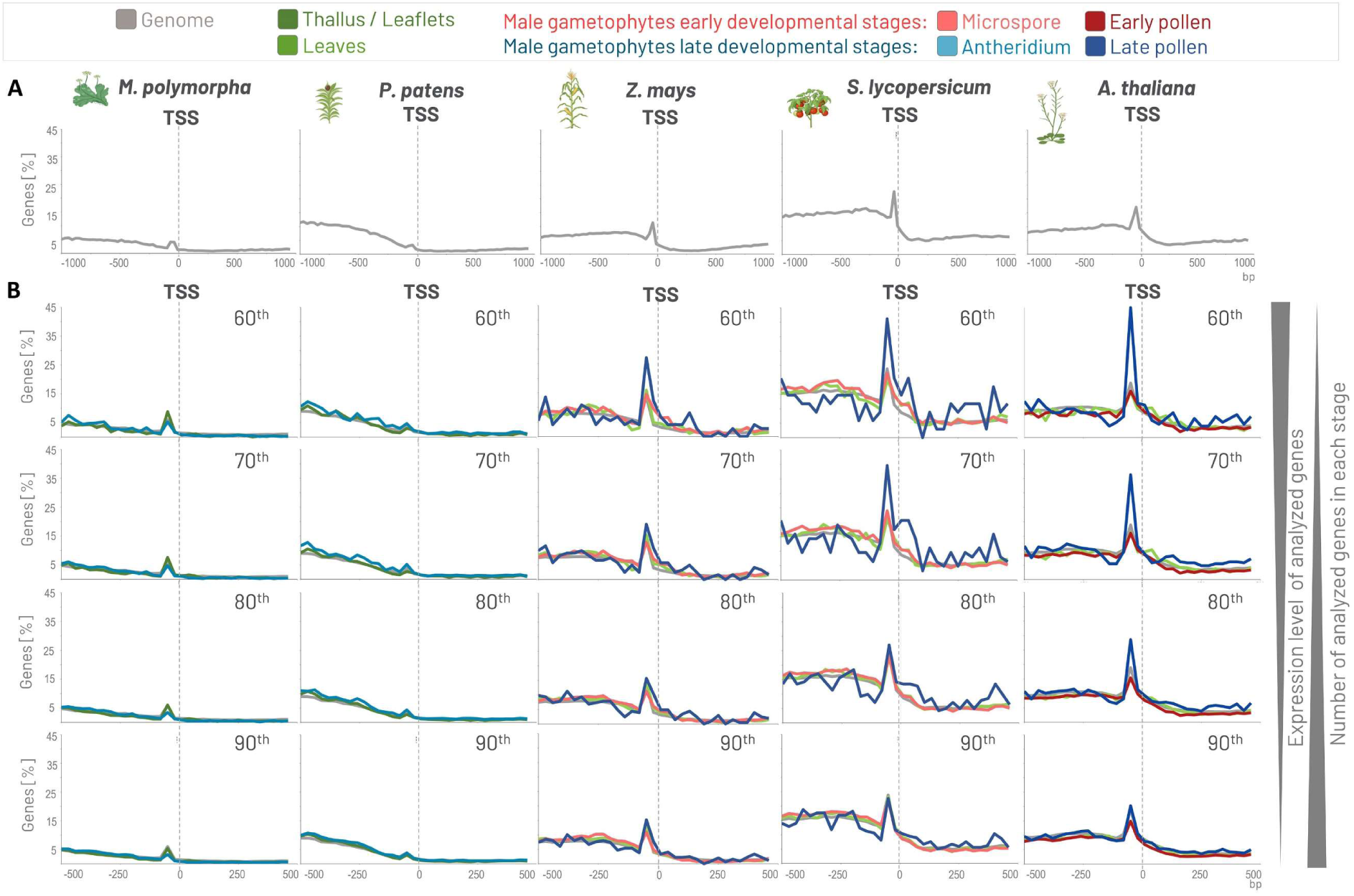
An example of the distribution of TATA-box in genomes across plant evolution and in genes expressed at various levels in male gametophyte tissues. (A) Distribution of the TATA-box shows that only a small fraction of plants genes actually harbors a TATA-box in their promoters in −30 to −25 area upstream of TSS, regardless of the transcription of those genes (genome). The motif was searched in the interval <−1000, 1000> bp from TSS within the bucket size 30 bp, and the axis size was adjusted to 45% in all species. (B) Genes whose transcripts represent 60%, 70%, 80% and 90% percent of all transcripts transcribed from the total number of protein-coding genes (60^th^, 70^th^, 80^th^ and 90^th^ percentile) were analyzed in various selected stages in *M. polymorpha*, *P. patens*, *Z. mays*, *S. lycopersicum* and *A. thaliana*. Genes highly transcribed during late male gametophyte development in flowering plants possess a higher percentage of the TATA-box motifs located upstream of TSS than genes transcribed during early pollen. The motif was searched in the interval <−500, 500> bp from TSS within the bucket size 30 bp, and the axis size was adjusted to 45% in all species.

Further, we analyzed the genes whose transcripts represent 60%, 70%, 80% and 90% percent of all transcripts transcribed from the total number of protein-coding genes (60^th^, 70^th^, 80^th^ and 90^th^ percentile) in various stages during the male gametophyte development and in the thallus/leaflets/leaves. Analysis of the promoters showed that TATA-box-containing promoters are associated with the genes expressed during the late pollen development, but not early pollen development, in flowering plants, especially in *S. lycopersicum* and *A. thaliana* (**Fig. 2B** **and** **Fig. 3**).

**Fig. 3.**
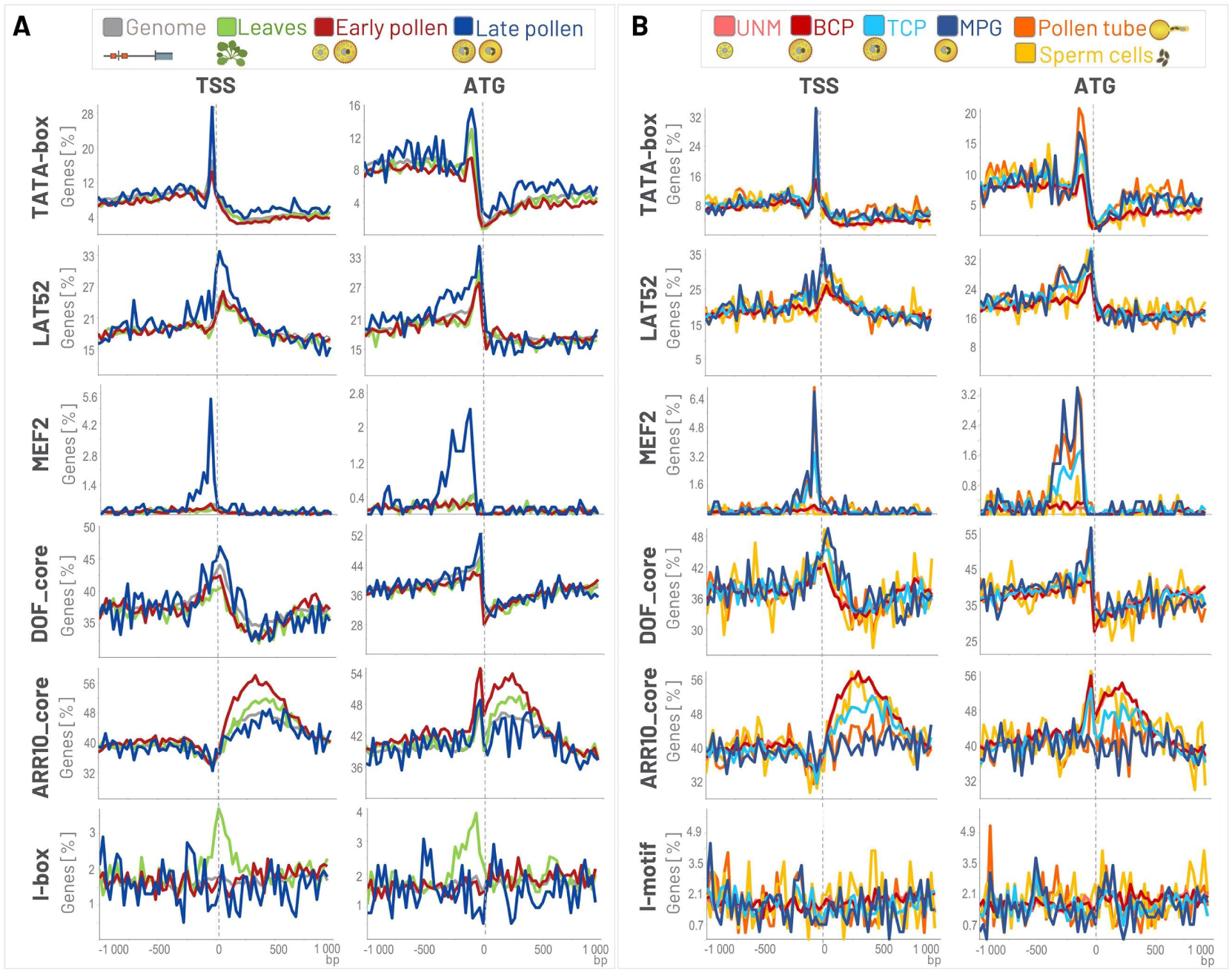
Example of the distribution of various motifs in the vicinity of TSS and ATG in *A. thaliana*. Colored lines represent different datasets and indicate the percentage of genes with the motifs at specific positions in the promoters of the genes whose transcripts represent 80% of all transcripts transcribed from the total number of protein-coding genes in each selected stage (80^th^ percentile): (A) in early pollen, late pollen, leaves, and regardless of the transcription level (genome); (B) in UNM, BCP, TCP, MPG, pollen tube and sperm cells. The motifs were searched in the interval <−1000, 1000> bp, within the bucket size 30 bp, and the axis size was adjusted. TATA-box (TATAWA); LAT52 (POLLEN1_LeLAT52, AGAAA); MEF2 (CTAWWWWTAG); DOF_core (AAAG); ARR10_core (GATY); I-motif (GATAAG); TSS, transcription start site; ATG, translation start site; UNM, uninucleate microspore; BCP, bicellular pollen; early pollen, UNM + BCP; TCP, tricellular pollen; MPG, mature pollen grain; late pollen, TCP + MPG; Pollen tube, semi-in vivo grown pollen tube; bp, base pair.

### GOLEM demonstrates distribution patterns of gene regulatory motifs linked to the male gametophyte

Unlike the position independence seen for many animal DNA sequence motifs, the activity of flowering plant DNA sequence motifs is strongly dependent on their position relative to the TSS [89]. Achieving precise localization and visualization of CREs regulatory motifs in genes that exhibit high transcription levels in various stages of male gametophyte development may help to elucidate the gene regulation in specific tissues.

LAT52 (also named POLLEN1_LeLAT52) is a pollen-specific motif (AGAAA) recognized within the promoter of *S. lycopersicum lat52* gene that encodes an essential protein expressed in the vegetative cell during pollen maturation. It specifically directs the transcription of genes required for PT growth and fertilization, ensuring successful reproduction in flowering plants [37,46]. Our analysis revealed that the preferential position of the LAT52 motif (the top of the peak) is located downstream of the TSS and upstream to ATG, i.e., in 5’UTR. Moreover, the number of the genes containing the LAT52 in *A. thaliana* is higher within the 5’UTR region of genes exhibiting elevated expression (80^th^ percentile) in late pollen development and PT, as opposed to genes expressed during early pollen development or in leaves, as was expected (**Fig. 3**).

MEF2-type CArG-box (CTA(A/T)_4_TAG) is bound by MADS-protein complexes functioning in mature pollen [36,90]. It was shown that MEF2-type boxes are strongly overrepresented in the proximal region of promoters that are activated during the last stages of pollen development [36]. Our analysis using the GOLEM program verified a strong overrepresentation of MEF2-type box in late pollen development in *A. thaliana* (**Fig. 3A**), especially in genes expressed in MPG and pollen tube, but not in sperm cells or UNM/BCP (**Fig. 3B**). The MEF2-type box present in the promoters of the genes expressed during the late pollen development is located −80 bp upstream TSS.

DOF motif is recognized by plant-specific DNA-binding TFs named Dof (DNA-binding with One Finger) domain proteins [38,39], which have crucial roles in many physiological processes, including hormone signaling and various biotic or abiotic stress responses but are also reported to regulate many biological processes, such as dormancy or tissue differentiation (reviewed in [60]). Using the program GOLEM, we have revealed the overrepresentation of DOF_core motif (AAAG) in genes activated during the last stages of pollen development (TCP, MPG) and in the sperm cell, compared to early pollen stages or leaves. Interestingly, the peak of DOF_core motif is centered −40 bp upstream of ATG in *A. thaliana*, predominantly located in the 5’UTR (**Fig. 3A, B**).

The distribution of the ARR10_core motif, which is recognized by the ARR10 protein. ARR10 is one of a type-B ARABIDOPSIS RESPONSE REGULATORS (B-ARRs) TFs that are associated with cytokinin transcriptional response network [27,51]. Cytokinins play a crucial role in regulating reproductive development in *Arabidopsis* (reviewed in [91]). Our analysis showed that the ARR10_core (GATY) motif is overrepresented in early pollen stages (UNM, BCP) and sperm cells, however, its presence is decreased in TCP and even more so in MPG stages in *A. thaliana*. The ARR10_core motifs are present not only in 5’UTRs but also within the gene bodies (**Fig. 3**).

### GOLEM disclose localization of gene regulatory motifs in sporophyte

Although our software, GOLEM, is primarily focused on tissues associated with male gametophyte development, it can also be utilized to search for gene regulatory motifs near the TSS and ATG in leaves, leaflets, and thallus of plant species available in GOLEM. Thus, it can visualize motifs associated with the regulation of genes involved not only in gametophyte development but also in sporophyte development in angiosperms.

The I-box has been suggested to be involved in light-regulated and/or leaf-specific gene expression of photosynthetic genes [29–31] and can be bound by myb-like proteins in *S. lycopersicum* [92]. The leaf-specific overrepresentation of I-box (GATAAG; **Supplementary Fig. 3**) can be tracked also using the GOLEM program. The I-box is overrepresented in the 5’UTR region of genes expressed in the sporophyte but not in the gametophyte, as detected not only in the exemplified *A. thaliana* (**Fig. 3**), but also in *S. lycopersicum* and *Z. mays* using the GOLEM program (data not shown - see in GOLEM program).

The DRE/CRT element is recognized by the drought-responsive element binding (DREB) family of TFs [62,83]. These cis-elements are located in promoter regions of target stress-responsive genes and play an important role in the regulation of stress-inducible transcription [61,63]. Therefore, it is not surprising that there are no significant changes in the overrepresentation of this element between sporophyte and gametophyte tissues that were not stressed, as expected. Interestingly, contrary to expectations, the DRE/CRT element is preferentially located in the gene bodies of genes that exhibit higher expression under non-stressed conditions, rather than in their promoters (**Supplementary Fig. 3**).

No significant changes in the overrepresentation between the gametophyte and sporophyte in *A. thaliana* were observed in other motifs present in the GOLEM software. For example, the BR-response element (CGTGYG) recognized by Brassinazole-resistant (BZR) family plant-specific TFs shows a negligible difference between sporophyte and gametophyte (**Supplementary Fig. 3**), even though BZRs play a significant role in regulating plant growth and development, as well as stress responses [57–59]. Similarly, the E-box (enhancer box, CANNTG), which is recognized by the helix-loop-helix (bHLH) family of TFs, and is important for plant growth, development, light signal transduction, and stress responses (reviewed in [52]), shows no difference between sporophyte and gametophyte.

### Further analysis of the genes with motif of interest in their promoters

The GOLEM program enables exporting normalized expression values of genes, depicted as TPM, from selected tissues at a specified percentile or for a chosen number of genes. This export also includes their expression levels in other tissues, formatted as a table in XLSX. Additionally, the table contains the gene identifier numbers of the genes that contain the motif of interest in a certain bucket. These gene accession numbers can be analyzed using various bioinformatical approaches, such as gene description search, gene ontology (GO) enrichment analysis, protein-protein interaction network analysis, or other relevant analyses.

To illustrate this feature, genes containing the LAT52 motif in their promoter were exported. LAT52 shows a sharp peak upstream of the ATG in genes expressed in the 80^th^ percentile during the late pollen stage in *A. thaliana* (**Fig. 3**, **Fig. 4A**). The XLSX table was exported from the late pollen stage using GOLEM, covering buckets between −70 to −10 bp from the ATG start codon (**Fig. 4B**). The gene identifier numbers (AGI - Arabidopsis Genome Initiative ID numbers) from this table, within the −70 to −10 bp region of the ATG start codon, were uploaded to the g:PROFILER for GO enrichment analysis. The GO analysis revealed that genes containing the LAT52 motif in the region −70 to −10 bp of their ATG are enriched in GO terms associated with biological processes (BP) such as pollination, PT growth and development, cytoskeleton organization, pectin catabolic processes, and mitochondrial ATP/ADP transport (**Fig. 4C**). All these terms are relevant to PT growth; for example, pectin plays a role in adhesion between the style and PT to prevent PT wandering, and cytoskeleton components like microtubules and actin filaments are involved in mitochondrial distribution in PT tip growth, as reviewed by [71].

**Fig. 4.**
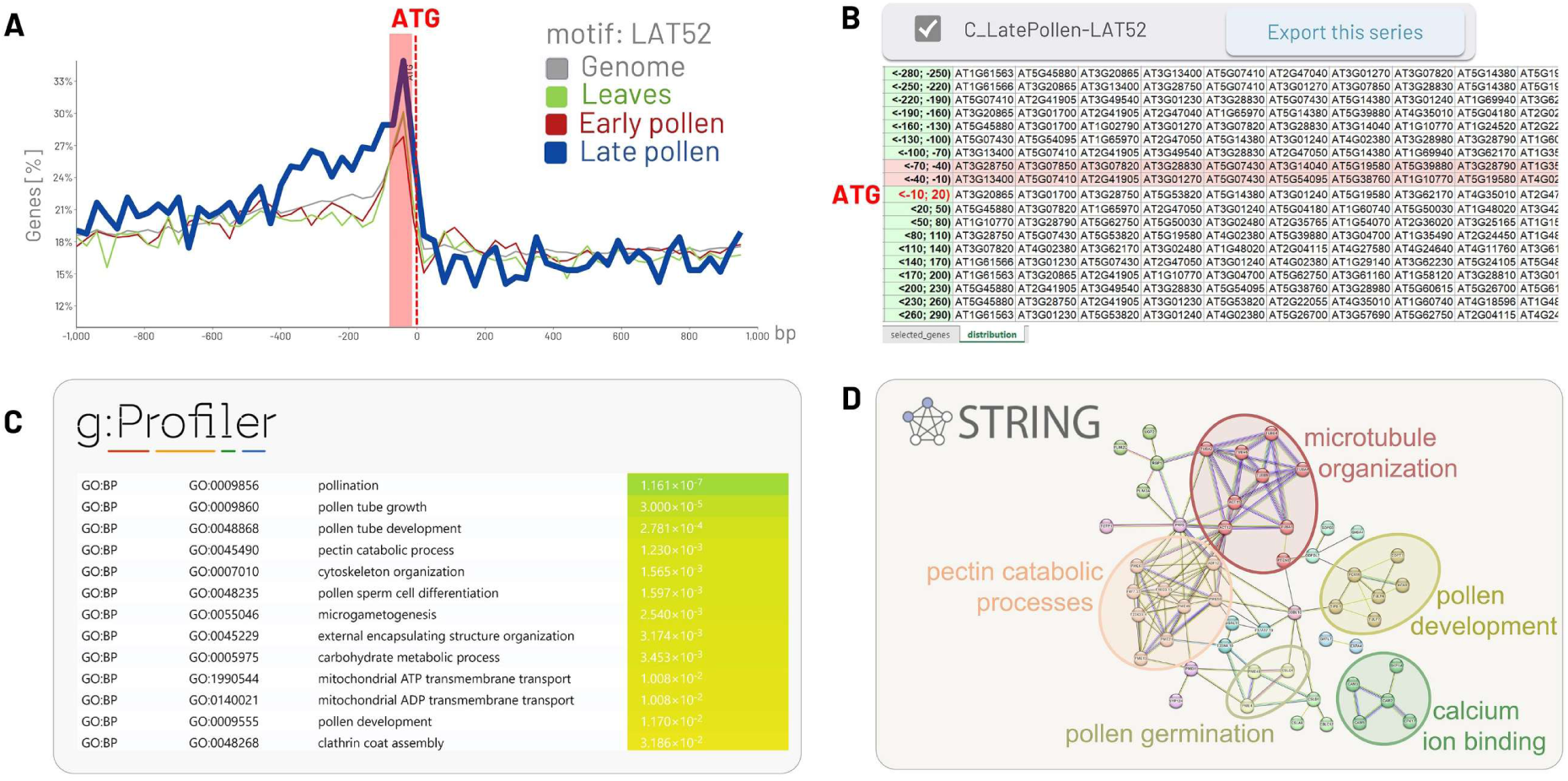
Functional analysis of the genes with LAT52 in the vicinity of ATG. (A) The genes expressed in the 80^th^ percentile during the late pollen stage in *A. thaliana*, <−1000, 1000> bp within the bucket size 30 bp were visualized using GOLEM software. (B) The XLSX table was exported from the late pollen stage using GOLEM. (C) The gene identifier numbers from XLSX table, covering buckets between <−70, −10) bp upstream from the ATG, were uploaded to the g:PROFILER for GO enrichment analysis. (D) The genes associated with the GO term biological processes (GO:BP) were uploaded to STRING to visualize the comprehensive network of protein-protein interactions of proteins whose genes are expressed during late pollen development and contain LAT52 upstream of ATG.

When genes associated with biological processes (BP) in g:PROFILER were uploaded to STRING, where a comprehensive network of predicted and known protein interactions was generated. These interactions, which include both physical and functional associations, revealed that genes containing the LAT52 motif within the region −70 to −10 bp upstream of the ATG, expressed during late pollen development in *A. thaliana*, are associated with biological processes such as pollen germination, pollen development, microtubule organization, pectin catabolic processes, and calcium ion binding (**Fig. 4D**). All these processes are crucial for PT growth, development, and male-female communication, as reviewed by [71].

## 4 Conclusion

Achieving accurate localization and visualization of gene regulatory motifs in promoters of genes with high transcription levels restricted to specific tissues involves a multi-step process. This process can be hindered by the user’s proficiency with various bioinformatics tools or the requirement for input data in specific formats. To address the challenge of precisely localizing and visualizing regulatory motifs near transcription and translation start sites - key elements in gene regulation in specific tissues - we introduced the GOLEM software. GOLEM provides a user-friendly platform for investigating the distribution of any motif of interest in gene promoters across diverse plant genomes and developmental tissues, with a particular emphasis on male gametophytes.

Using gene regulatory motifs such as LAT52, MEF2, DOF_core, and ARR10_core - previously implicated in the regulation of pollen and/or plant development - we demonstrated that the GOLEM program is an effective tool for visualizing the distribution of these motifs within gene promoters. Our analysis with GOLEM revealed whether these motifs are preferentially associated with genes expressed during the early or late stages of male gametophyte development. Additionally, GOLEM enables accuratelly map the positional distribution of peaks near the TSS or ATG, even within the 5’ UTR, across various species. Beyond gametophyte-specific motifs, GOLEM also facilitates the visualization of motifs in plant sporophytes (e.g., leaves). For instance, GOLEM has shown that I-box motifs, which are associated with plant photosynthesis, are overrepresented in the 5’ UTR region of genes expressed in the sporophyte but not in the gametophyte. Furthermore, GOLEM allows users to track all analyses and export data on genes with motifs of interest at specific positions relative to the TSS/ATG for further analysis using tools such as Gene Ontology (GO) or STRING.

Overall, the user-friendly online software GOLEM is a valuable resource for elucidating the abundance, distribution, and tissue-specific association of any motif of interest across diverse plant species and evolutionary stages. As GOLEM does not require programming skills or advanced bioinformatics expertise, it is particularly well-suited for biologists with limited experience in complex bioinformatics tools.

## Acknowledgement

Biological Data Management and Analysis Core Facility of CEITEC Masaryk University, funded by ELIXIR CZ research infrastructure (MEYS Grant No: LM2023055), is gratefully acknowledged for supporting the research presented in this paper. We also extend our gratitude to Mgr. Jiří Rudolf for his valuable comments and fruitful discussions.

## Author contribution

P.P.S. and D.H. conceived the study. B.K. analyzed RNA-seq data, calculated the TPM. L.N. imposed computation analysis and visualization tool. T.R. and R.S. implemented the website and supported program accessibility. T.P. and A.K. helped with analysis of exported data. P.P.S wrote the paper with the help of all co-authors.

## Data availability

Availability and implementation: GOLEM is freely available at https://golem.ncbr.muni.cz and its source codes are provided under the MIT licence at GitHub at https://github.com/sb-ncbr/golem.

## Supplementary Material

### Figures

**Supplementary Fig. 1.**
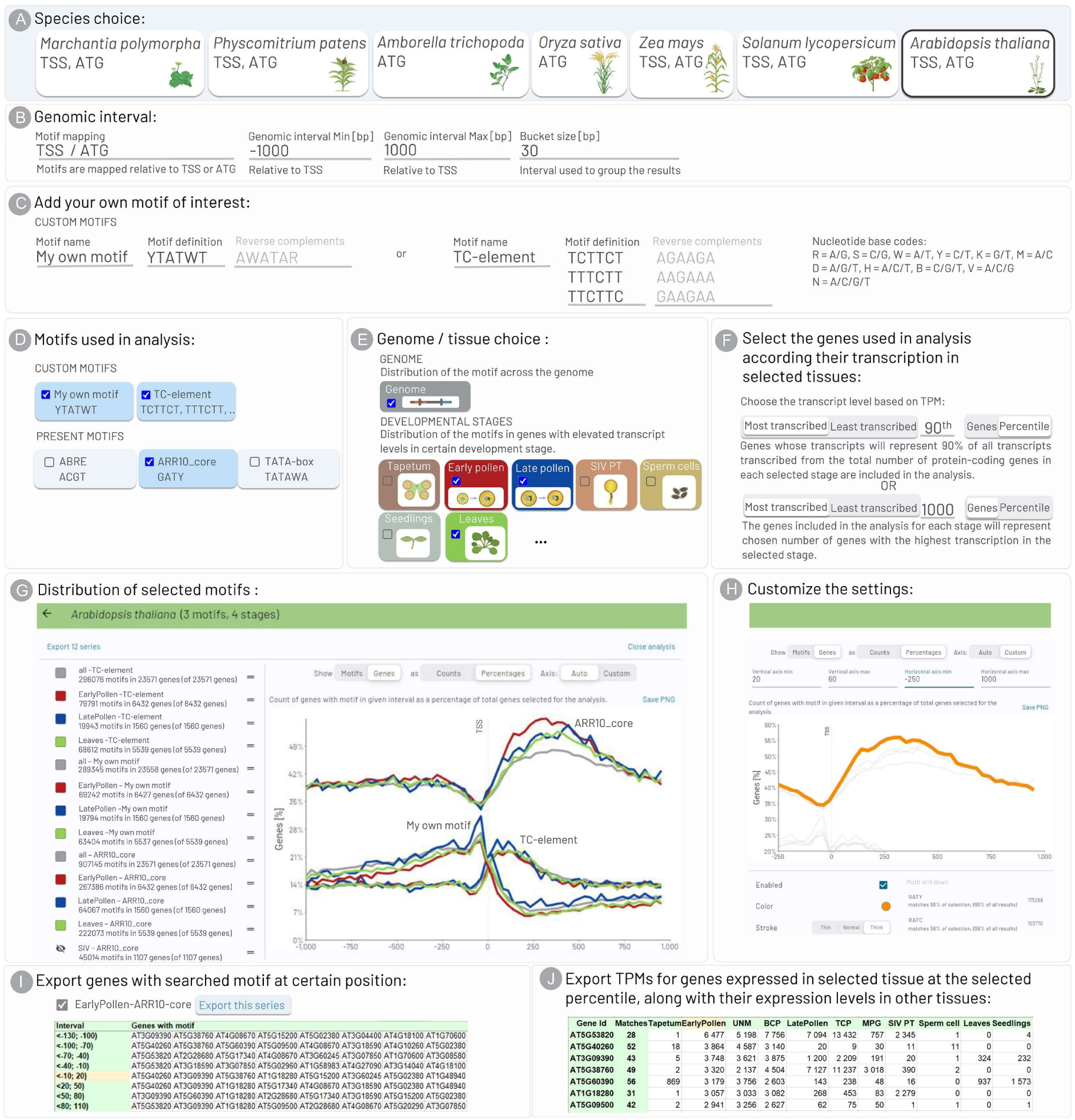
A detailed overview of the workflow of the GOLEM software. (A) One plant species across the plant Tree of Life is chosen, and the data are downloaded on the web browser (*Marchantia*, *Physcomitrium*, *Amborella*, *Oryza*, *Zea*, *Solanum*, and *Arabidopsis*). If available, the positions of both TSS and ATG are given. (B) The defined region (genomic interval) in the vicinity of the TSS or ATG, within the selected bucket size (bp), is chosen. (C) A single custom motif of interest can be defined by users. Additionally, multiple motifs as well as degenerate motifs can also be searched for by users. (D) Optionally the motif can be chosen from several motifs present in the software. (E) The promoters of genes showing expression in selected tissues and developmental stages (sporophyte, male gametophyte), along with an analysis of genome-wide distribution regardless of transcription, are chosen for the analysis. (F) The selection of genes that are highly or minimally transcribed in tissues or developmental stages of interest can be determined by the user, based on a specified percentile (default is the 90^th^ percentile) or a certain number of genes included in the analysis. (G) The exemplified “My own motif, TC-element and ARR10_core” motifs show various distributions upstream/downstream of TSS. The motif TC-element shows higher prevalence in the promoters of genes transcribed during Late pollen development (blue) and the motif ARR10_core shows higher prevalence in the promoters of the genes transcribed during early pollen (red) stages, in comparison to the genome-wide distribution (all; grey). The symbol (=) is used to change the curve order. The individual stages can be made invisible. (H) The customization options for the output graph include adjusting curve color/stroke, axes size, and displaying either percentages or counts of genes with the motif of interest. Additionally, the output graph can be saved in PNG format. (I) The accession numbers of genes with certain motif at the selected interval may be exported in XLSX format tables. (J) The normalized expression values of genes, represented as Transcript Per Million (TPM) in selected tissue at a specified percentile or for a chosen number of genes, along with their expression in other tissues, can be exported as a table in XLSX format. Plant icons were created with BioRender.com.

**Supplementary Fig. 2.**
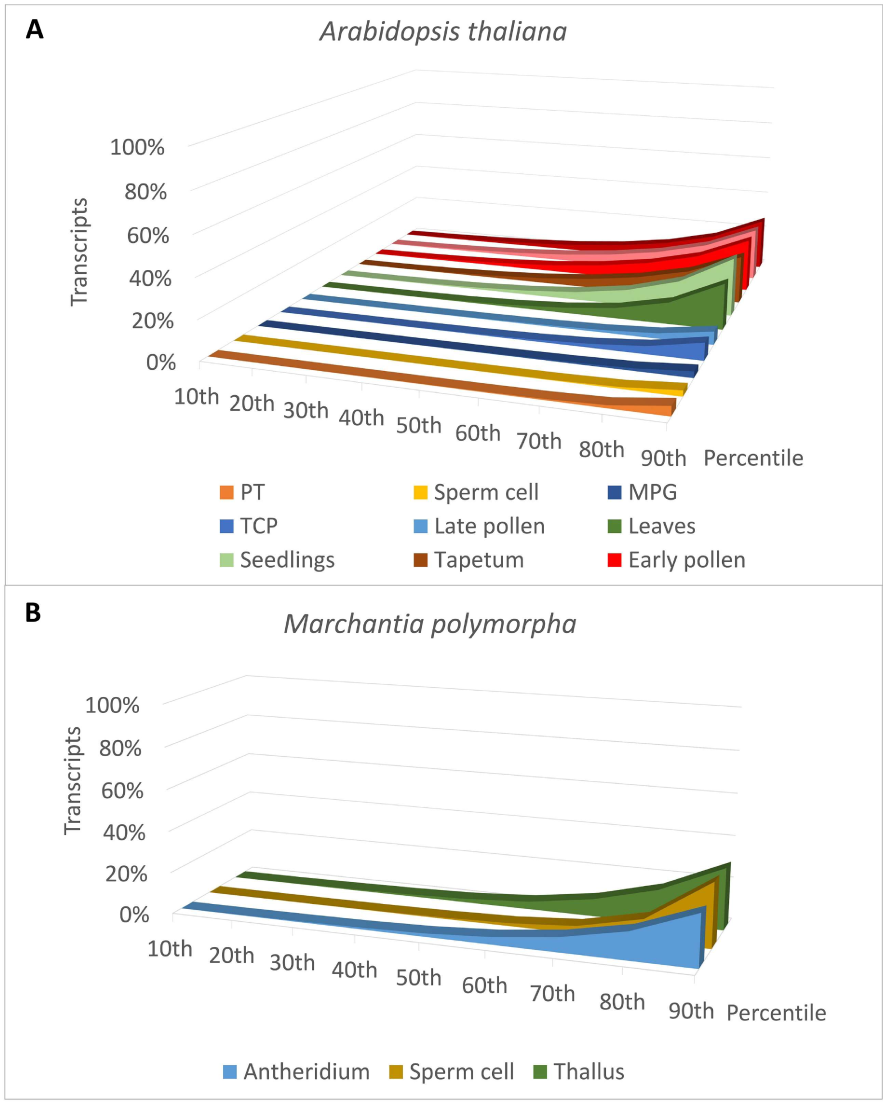
The number of genes contributing to expression programs varies between developmental stages or tissues. (A) In early pollen stages, leaves and seedlings of *A. thaliana* the genes whose transcripts account for 90% of all transcripts transcribed from the total number of protein-coding genes (90^th^ percentile) represent 27%, 24% and 30% of the total protein-coding genes, respectively. In late pollen stages, sperm cells and PT, those genes represent 7%, 3% and 5%, respectively. (B) In bryophyte *M. polymorpha*, the genes whose transcripts comprise 90^th^ percentile show more similar levels in antheridia, sperm cells and thallus, 25%, 30% and 29%, respectively. UNM, uninucleate microspore; BCP, bicellular pollen; early pollen, UNM + BCP; TCP, tricellular pollen; MPG, mature pollen grain; late pollen, TCP + MPG; PT, semi-in vivo grown pollen tube; Percentile, genes whose transcripts represent certain percent of all transcripts transcribed from the total number of protein-coding genes in each selected stage.

**Supplementary Fig. 3.**
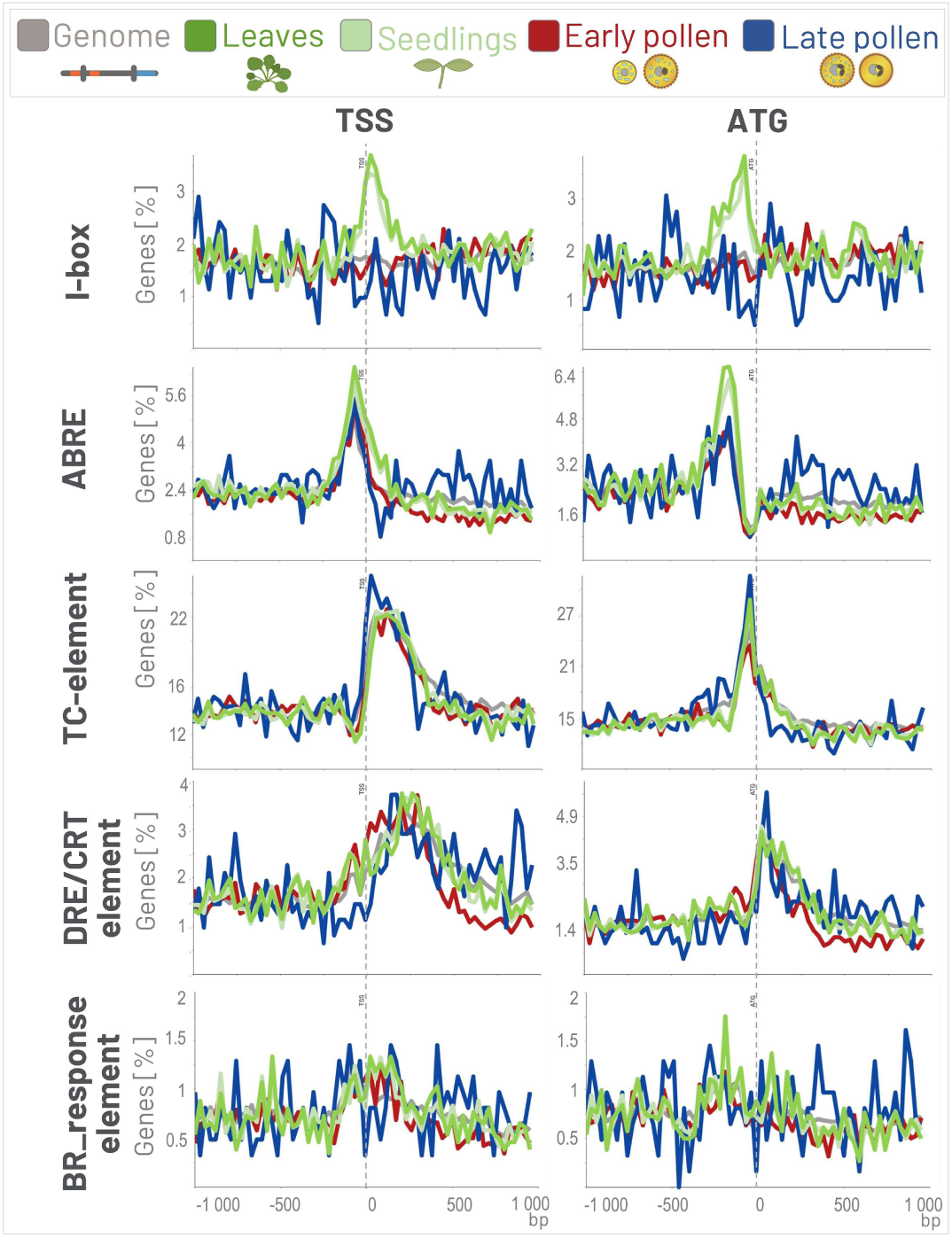
Example of the distribution of various motifs in the vicinity of TSS and ATG in *A. thaliana* with a focus on plant leaves and seedlings. Colored lines represent different datasets and indicate the percentage of genes containing selected motifs at specific positions in the promoters of the genes whose transcripts represent 80% of all transcripts transcribed from the total number of protein-coding genes in each selected stage: early pollen, late pollen, leaves, seedling and regardless of the transcription level (genome). The motifs were searched in the interval <−1000, 1000> bp, within the bucket size 30 bp, and the axis size was adjusted. I-motif (GATAAG); ABRE (ACGTG); TC_element (TCTTCT, TTTCTT, TTCTTC); DRE/CRT_element (CANNTG); BR_response element (CGTGYG); TSS, transcription start site; ATG, translation start site; early pollen, UNM + BCP; late pollen, TCP + MPG; bp, base pair.

### Tables

**Supplementary Table 1.** (A) The reference genomes and genome annotations files used in GOLEM software. (B) The tissues/male gametophyte developmental stages present in GOLEM software, along with the source of Transcript Per Million (TPM) values or RNA-seq datasets used for their calculation.

**Supplementary Table 2.** Normalized TPM values used for the data processing pipeline.

## References

1. Galli M, Feng F, Gallavotti A. Mapping Regulatory Determinants in Plants. *Front Genet*. Frontiers; 2020; doi: 10.3389/fgene.2020.591194.

2. Schmitz RJ, Grotewold E, Stam M. Cis-regulatory sequences in plants: Their importance, discovery, and future challenges. Plant Cell. 2022; doi: 10.1093/plcell/koab281.

3. Preissl S, Gaulton KJ, Ren B. Characterizing cis-regulatory elements using single-cell epigenomics. Nat Rev Genet. Nature Publishing Group; 2023; doi: 10.1038/s41576-022-00509-1.

4. Bailey TL, Johnson J, Grant CE, Noble WS. The MEME Suite. Nucleic Acids Res. 2015; doi: 10.1093/nar/gkv416.

5. Grant CE, Bailey TL, Noble WS. FIMO: scanning for occurrences of a given motif. Bioinformatics. 2011; doi: 10.1093/bioinformatics/btr064.

6. Blaby IK, Blaby-Haas CE, Tourasse N, Hom EFY, Lopez D, Aksoy M, et al.. The Chlamydomonas genome project: a decade on. Trends Plant Sci. 2014; doi: 10.1016/j.tplants.2014.05.008.

7. McCouch SR, CGSNL (Committee on Gene Symbolization N and L Rice Genetics Cooperative). Gene Nomenclature System for Rice. Rice. 2008; doi: 10.1007/s12284-008-9004-9.

8. Pan R, Hu H, Xiao Y, Xu L, Xu Y, Ouyang K, et al. High-quality wild barley genome assemblies and annotation with Nanopore long reads and Hi-C sequencing data. Sci Data. Nature Publishing Group; 2023; doi: 10.1038/s41597-023-02434-2.

9. Rensing SA, Goffinet B, Meyberg R, Wu S-Z, Bezanilla M. The Moss *Physcomitrium* (*Physcomitrella*) *patens*: A Model Organism for Non-Seed Plants. Plant Cell. 2020; doi: 10.1105/tpc.19.00828.

10. Tyagi P, Singh D, Mathur S, Singh A, Ranjan R. Upcoming progress of transcriptomics studies on plants: An overview. *Front Plant Sci*. Frontiers; 2022; doi: 10.3389/fpls.2022.1030890.

11. Julca I, Ferrari C, Flores-Tornero M, Proost S, Lindner A-C, Hackenberg D, et al. Comparative transcriptomic analysis reveals conserved programmes underpinning organogenesis and reproduction in land plants. Nat Plants. Nature Publishing Group; 2021; doi: 10.1038/s41477-021-00958-2.

12. Klodová B, Potěšil D, Steinbachová L, Michailidis C, Lindner A-C, Hackenberg D, et al. Regulatory dynamics of gene expression in the developing male gametophyte of *Arabidopsis*. Plant Reprod. 2023; doi: 10.1007/s00497-022-00452-5.

13. Shi D, Jouannet V, Agustí J, Kaul V, Levitsky V, Sanchez P, et al. Tissue-specific transcriptome profiling of the *Arabidopsis* inflorescence stem reveals local cellular signatures. Plant Cell. 2021; doi: 10.1093/plcell/koaa019.

14. Feng Y, Zhang Y, Ebright RH. Structural basis of transcription activation. Science. 2016; doi: 10.1126/science.aaf4417.

15. Lifton RP, Goldberg ML, Karp RW, Hogness DS. The organization of the histone genes in *Drosophila melanogaster*: functional and evolutionary implications. Cold Spring Harb Symp Quant Biol. 1978; doi: 10.1101/sqb.1978.042.01.105.

16. Suzuki Y, Tsunoda T, Sese J, Taira H, Mizushima-Sugano J, Hata H, et al. Identification and characterization of the potential promoter regions of 1031 kinds of human genes. Genome Res. 2001; doi: 10.1101/gr.gr-1640r.

17. McKnight SL, Kingsbury R. Transcriptional Control Signals of a Eukaryotic Protein-Coding Gene. Science. American Association for the Advancement of Science; 1982; doi: 10.1126/science.6283634.

18. Basehoar AD, Zanton SJ, Pugh BF. Identification and distinct regulation of yeast TATA box-containing genes. Cell. 2004; doi: 10.1016/s0092-8674(04)00205-3.

19. Yang C, Bolotin E, Jiang T, Sladek FM, Martinez E. Prevalence of the initiator over the TATA box in human and yeast genes and identification of DNA motifs enriched in human TATA-less core promoters. Gene. 2007; doi: 10.1016/j.gene.2006.09.029.

20. Carninci P, Sandelin A, Lenhard B, Katayama S, Shimokawa K, Ponjavic J, et al. Genome-wide analysis of mammalian promoter architecture and evolution. Nat Genet. Nature Publishing Group; 2006; doi: 10.1038/ng1789.

21. Shi W, Zhou W. Frequency distribution of TATA Box and extension sequences on human promoters. BMC Bioinformatics. 2006; doi: 10.1186/1471-2105-7-S4-S2.

22. Bernard V, Brunaud V, Lecharny A. TC-motifs at the TATA-box expected position in plant genes: a novel class of motifs involved in the transcription regulation. BMC Genomics. 2010; doi: 10.1186/1471-2164-11-166.

23. Savinkova LK, Sharypova EB, Kolchanov NA. On the Role of TATA Boxes and TATA-Binding Protein in *Arabidopsis thaliana*. Plants. Multidisciplinary Digital Publishing Institute; 2023; doi: 10.3390/plants12051000.

24. Civán P, Svec M. Genome-wide analysis of rice (*Oryza sativa L.* subsp. *japonica*) TATA box and Y Patch promoter elements. Genome. 2009; doi: 10.1139/G09-001.

25. Pavlu S, Nikumbh S, Kovacik M, An T, Lenhard B, Simkova H, et al. Core promoterome of barley embryo. Comput Struct Biotechnol J. 2024; doi: 10.1016/j.csbj.2023.12.003.

26. Vo Ngoc L, Wang Y-L, Kassavetis GA, Kadonaga JT. The punctilious RNA polymerase II core promoter. Genes Dev. 2017; doi: 10.1101/gad.303149.117.

27. Hosoda K, Imamura A, Katoh E, Hatta T, Tachiki M, Yamada H, et al. Molecular structure of the GARP family of plant Myb-related DNA binding motifs of the *Arabidopsis* response regulators. Plant Cell. 2002; doi: 10.1105/tpc.002733.

28. Šmeringai J, Schrumpfová PP, Pernisová M. Cytokinins – regulators of de novo shoot organogenesis. *Front Plant Sci*. Frontiers; 2023; doi: 10.3389/fpls.2023.1239133.

29. Castresana C, Staneloni R, Malik VS, Cashmore AR. Molecular characterization of two clusters of genes encoding the Type I CAB polypeptides of PSII in *Nicotiana plumbaginifolia*. Plant Mol Biol. 1987; doi: 10.1007/BF00016149.

30. Gidoni D, Brosio P, Bond-Nutter D, Bedbrook J, Dunsmuir P. Novel cis-acting elements in petunia Cab gene promoters. Mol Gen Genet MGG. 1989; doi: 10.1007/BF00339739.

31. Manzara T, Carrasco P, Gruissem W. Developmental and organ-specific changes in promoter DNA-protein interactions in the tomato rbcS gene family. Plant Cell. 1991; doi: 10.1105/tpc.3.12.1305.

32. She W, Baroux C. Chromatin dynamics during plant sexual reproduction. *Front Plant Sci*. Frontiers; 2014; doi: 10.3389/fpls.2014.00354.

33. Meissner ST. Plant sexual reproduction: perhaps the current plant two-sex model should be replaced with three- and four-sex models? Plant Reprod. 2021; doi: 10.1007/s00497-021-00420-5.

34. Williams JH, Reese JB. Chapter Twelve - Evolution of development of pollen performance. In: Grossniklaus U, editor. Curr Top Dev Biol. Academic Press; 2019; 10.1016/bs.ctdb.2018.11.012

35. Marand AP, Eveland AL, Kaufmann K, Springer NM. cis-Regulatory Elements in Plant Development, Adaptation, and Evolution. Annu Rev Plant Biol. Annual Reviews; 2023; doi: 10.1146/annurev-arplant-070122-030236.

36. Verelst W, Saedler H, Münster T. MIKC* MADS-Protein Complexes Bind Motifs Enriched in the Proximal Region of Late Pollen-Specific *Arabidopsis* Promoters. Plant Physiol. 2007; doi: 10.1104/pp.106.089805.

37. Bate N, Twell D. Functional architecture of a late pollen promoter: pollen-specific transcription is developmentally regulated by multiple stage-specific and co-dependent activator elements. Plant Mol Biol. 1998; doi: 10.1023/A:1006095023050.

38. Li J, Yuan J, Li M. Characterization of Putative cis-Regulatory Elements in Genes Preferentially Expressed in *Arabidopsis* Male Meiocytes. BioMed Res Int. 2014; doi: 10.1155/2014/708364.

39. Yanagisawa S. The Dof family of plant transcription factors. Trends Plant Sci. 2002; doi: 10.1016/S1360-1385(02)02362-2.

40. Hoffmann RD, Olsen LI, Husum JO, Nicolet JS, Thøfner JFB, Wätjen AP, et al. A *cis*-Regulatory Sequence Acts as a Repressor in the *Arabidopsis thaliana* Sporophyte but as an Activator in Pollen. Mol Plant. 2017; doi: 10.1016/j.molp.2016.12.010.

41. Peters B, Casey J, Aidley J, Zohrab S, Borg M, Twell D, et al. A Conserved cis-Regulatory Module Determines Germline Fate through Activation of the Transcription Factor DUO1 Promoter. Plant Physiol. 2017; doi: 10.1104/pp.16.01192.

42. Sharma N, Russell SD, Bhalla PL, Singh MB. Putative cis-regulatory elements in genes highly expressed in rice sperm cells. BMC Res Notes. 2011; doi: 10.1186/1756-0500-4-319.

43. Dobin A, Davis CA, Schlesinger F, Drenkow J, Zaleski C, Jha S, et al. STAR: ultrafast universal RNA-seq aligner. Bioinformatics. 2013; doi: 10.1093/bioinformatics/bts635.

44. Li B, Dewey CN. RSEM: accurate transcript quantification from RNA-Seq data with or without a reference genome. BMC Bioinformatics. 2011; doi: 10.1186/1471-2105-12-323.

45. Hamilton DA, Schwarz YH, Mascarenhas JP. A monocot pollen-specific promoter contains separable pollen-specific and quantitative elements. Plant Mol Biol. 1998; doi: 10.1023/A:1006083725102.

46. Muschietti J, Dircks L, Vancanneyt G, McCormick S. LAT52 protein is essential for tomato pollen development: pollen expressing antisense LAT52 RNA hydrates and germinates abnormally and cannot achieve fertilization. Plant J. 1994; doi: 10.1046/j.1365-313X.1994.06030321.x.

47. Peng J, Qi X, Chen X, Li N, Yu J. ZmDof30 Negatively Regulates the Promoter Activity of the Pollen-Specific Gene Zm908. *Front Plant Sci*. Frontiers; 2017; doi: 10.3389/fpls.2017.00685.

48. Rogers HJ, Bate N, Combe J, Sullivan J, Sweetman J, Swan C, et al. Functional analysis of cis-regulatory elements within the promoter of the tobacco late pollen gene g10. Plant Mol Biol. 2001; doi: 10.1023/A:1010695226241.

49. Hattori T, Totsuka M, Hobo T, Kagaya Y, Yamamoto-Toyoda A. Experimentally Determined Sequence Requirement of ACGT-Containing Abscisic Acid Response Element. Plant Cell Physiol. 2002; doi: 10.1093/pcp/pcf014.

50. Watanabe KA, Homayouni A, Gu L, Huang K-Y, Ho T-HD, Shen QJ. Transcriptomic analysis of rice aleurone cells identified a novel abscisic acid response element. Plant Cell Environ. 2017; doi: 10.1111/pce.13006.

51. Xie M, Chen H, Huang L, O’Neil RC, Shokhirev MN, Ecker JR. A B-ARR-mediated cytokinin transcriptional network directs hormone cross-regulation and shoot development. Nat Commun. Nature Publishing Group; 2018; doi: 10.1038/s41467-018-03921-6.

52. Hao Y, Zong X, Ren P, Qian Y, Fu A. Basic Helix-Loop-Helix (bHLH) Transcription Factors Regulate a Wide Range of Functions in *Arabidopsis*. Int J Mol Sci. Multidisciplinary Digital Publishing Institute; 2021; doi: 10.3390/ijms22137152.

53. Shen Q, Ho TH. Functional dissection of an abscisic acid (ABA)-inducible gene reveals two independent ABA-responsive complexes each containing a G-box and a novel cis-acting element. Plant Cell. 1995; doi: 10.1105/tpc.7.3.295.

54. Yamaguchi-Shinozaki K, Mundy J, Chua NH. Four tightly linked rab genes are differentially expressed in rice. Plant Mol Biol. 1990; doi: 10.1007/BF00015652.

55. Ohme-Takagi M, Shinshi H. Ethylene-inducible DNA binding proteins that interact with an ethylene-responsive element. Plant Cell. 1995; doi: 10.1105/tpc.7.2.173.

56. Zhang H, Xie B, Lu X, Yang Y, Huang R. GCC box inArabidopsis PDF1.2 promoter is an essential and sufficient cis-acting element in response to MeJA treatment. Chin Sci Bull. 2004; doi: 10.1007/BF03183717.

57. Chen X, Wu X, Qiu S, Zheng H, Lu Y, Peng J, et al. Genome-Wide Identification and Expression Profiling of the BZR Transcription Factor Gene Family in *Nicotiana benthamiana*. Int J Mol Sci. Multidisciplinary Digital Publishing Institute; 2021; doi: 10.3390/ijms221910379.

58. Nolan TM, Vukašinović N, Liu D, Russinova E, Yin Y. Brassinosteroids: Multidimensional Regulators of Plant Growth, Development, and Stress Responses[OPEN]. Plant Cell. 2020; doi: 10.1105/tpc.19.00335.

59. Wang Z-Y, Nakano T, Gendron J, He J, Chen M, Vafeados D, et al. Nuclear-Localized BZR1 Mediates Brassinosteroid-Induced Growth and Feedback Suppression of Brassinosteroid Biosynthesis. Dev Cell. 2002; doi: 10.1016/S1534-5807(02)00153-3.

60. Zou X, Sun H. DOF transcription factors: Specific regulators of plant biological processes. *Front Plant Sci*. Frontiers; 2023; doi: 10.3389/fpls.2023.1044918.

61. Agarwal PK, Gupta K, Lopato S, Agarwal P. Dehydration responsive element binding transcription factors and their applications for the engineering of stress tolerance. J Exp Bot. 2017; doi: 10.1093/jxb/erx118.

62. Yamaguchi-Shinozaki K, Shinozaki K. A novel cis-acting element in an *Arabidopsis* gene is involved in responsiveness to drought, low-temperature, or high-salt stress. Plant Cell. 1994; doi: 10.1105/tpc.6.2.251.

63. Yang Y, Al-Baidhani HHJ, Harris J, Riboni M, Li Y, Mazonka I, et al. DREB/CBF expression in wheat and barley using the stress-inducible promoters of HD-Zip I genes: impact on plant development, stress tolerance and yield. Plant Biotechnol J. 2020; doi: 10.1111/pbi.13252.

64. Meiller D. Modern App Development with Dart and Flutter 2: A Comprehensive Introduction to Flutter. Walter de Gruyter GmbH & Co KG; 2021; ISBN-10 3110721279

65. Kolberg L, Raudvere U, Kuzmin I, Adler P, Vilo J, Peterson H. g:Profiler—interoperable web service for functional enrichment analysis and gene identifier mapping (2023 update). Nucleic Acids Res. 2023; doi: 10.1093/nar/gkad347.

66. Reimand J, Kull M, Peterson H, Hansen J, Vilo J. g:Profiler—a web-based toolset for functional profiling of gene lists from large-scale experiments. Nucleic Acids Res. 2007; doi: 10.1093/nar/gkm226.

67. Snel B, Lehmann G, Bork P, Huynen MA. STRING: a web-server to retrieve and display the repeatedly occurring neighbourhood of a gene. Nucleic Acids Res. 2000; doi: 10.1093/nar/28.18.3442.

68. Szklarczyk D, Kirsch R, Koutrouli M, Nastou K, Mehryary F, Hachilif R, et al. The STRING database in 2023: protein-protein association networks and functional enrichment analyses for any sequenced genome of interest. Nucleic Acids Res. 2023; doi: 10.1093/nar/gkac1000.

69. Bokvaj P, Hafidh S, Honys D. Transcriptome profiling of male gametophyte development in *Nicotiana tabacum*. Genomics Data. 2015; doi: 10.1016/j.gdata.2014.12.002.

70. Hafidh S, Fíla J, Honys D. Male gametophyte development and function in angiosperms: a general concept. Plant Reprod. 2016; doi: 10.1007/s00497-015-0272-4.

71. Hafidh S, Honys D. Reproduction Multitasking: The Male Gametophyte. Annu Rev Plant Biol. Annual Reviews; 2021; doi: 10.1146/annurev-arplant-080620-021907.

72. Williams JH, Taylor ML, O’Meara BC. Repeated evolution of tricellular (and bicellular) pollen. Am J Bot. 2014; doi: 10.3732/ajb.1300423.

73. Johnson MA, Harper JF, Palanivelu R. A Fruitful Journey: Pollen Tube Navigation from Germination to Fertilization. Annu Rev Plant Biol. Annual Reviews; 2019; doi: 10.1146/annurev-arplant-050718-100133.

74. Kohchi T, Yamato KT, Ishizaki K, Yamaoka S, Nishihama R. Development and Molecular Genetics of *Marchantia polymorpha*. Annu Rev Plant Biol. Annual Reviews; 2021; doi: 10.1146/annurev-arplant-082520-094256.

75. Hafidh S, Potě¡il D, Müller K, Fíla J, Michailidis C, Herrmannová A, et al. Dynamics of the Pollen Sequestrome Defined by Subcellular Coupled Omics. Plant Physiol. 2018; doi: 10.1104/pp.18.00648.

76. Zhao S, Ye Z, Stanton R. Misuse of RPKM or TPM normalization when comparing across samples and sequencing protocols. RNA N Y N. 2020; doi: 10.1261/rna.074922.120.

77. Honys D, Twell D. Comparative Analysis of the *Arabidopsis* Pollen Transcriptome. Plant Physiol. 2003; doi: 10.1104/pp.103.020925.

78. Honys D, Twell D. Transcriptome analysis of haploid male gametophyte development in *Arabidopsis*. Genome Biol. 2004; doi: 10.1186/gb-2004-5-11-r85.

79. Bowman JL, Kohchi T, Yamato KT, Jenkins J, Shu S, Ishizaki K, et al. Insights into Land Plant Evolution Garnered from the *Marchantia polymorpha* Genome. Cell. 2017; doi: 10.1016/j.cell.2017.09.030.

80. Zhang H, Wang Y, Wu X, Tang X, Wu C, Lu J. Determinants of genome-wide distribution and evolution of uORFs in eukaryotes. Nat Commun. Nature Publishing Group; 2021; doi: 10.1038/s41467-021-21394-y.

81. Srivastava AK, Lu Y, Zinta G, Lang Z, Zhu J-K. UTR-Dependent Control of Gene Expression in Plants. Trends Plant Sci. 2018; doi: 10.1016/j.tplants.2017.11.003.

82. Yu C-P, Lin J-J, Li W-H. Positional distribution of transcription factor binding sites in *Arabidopsis thaliana*. Sci Rep. Nature Publishing Group; 2016; doi: 10.1038/srep25164.

83. Champ KI, Febres VJ, Moore GA. The role of CBF transcriptional activators in two Citrus species (Poncirus and Citrus) with contrasting levels of freezing tolerance. Physiol Plant. 2007; doi: 10.1111/j.1399-3054.2006.00826.x.

84. Boyce JM, Knight H, Deyholos M, Openshaw MR, Galbraith DW, Warren G, et al. The sfr6 mutant of *Arabidopsis* is defective in transcriptional activation via CBF/DREB1 and DREB2 and shows sensitivity to osmotic stress. Plant J. 2003; doi: 10.1046/j.1365-313X.2003.01734.x.

85. Knight H, Mugford SG, Ülker B, Gao D, Thorlby G, Knight MR. Identification of SFR6, a key component in cold acclimation acting post-translationally on CBF function. Plant J. 2009; doi: 10.1111/j.1365-313X.2008.03763.x.

86. Liu Q, Kasuga M, Sakuma Y, Abe H, Miura S, Yamaguchi-Shinozaki K, et al. Two Transcription Factors, DREB1 and DREB2, with an EREBP/AP2 DNA Binding Domain Separate Two Cellular Signal Transduction Pathways in Drought- and Low-Temperature-Responsive Gene Expression, Respectively, in *Arabidopsis*. Plant Cell. 1998; doi: 10.1105/tpc.10.8.1391.

87. Vazquez-Hernandez M, Romero I, Escribano MI, Merodio C, Sanchez-Ballesta MT. Deciphering the Role of CBF/DREB Transcription Factors and Dehydrins in Maintaining the Quality of Table Grapes cv. Autumn Royal Treated with High CO2 Levels and Stored at 0°C. Front Plant Sci. Frontiers; 2017; doi: 10.3389/fpls.2017.01591.

88. Shiu S-H, Shih M-C, Li W-H. Transcription Factor Families Have Much Higher Expansion Rates in Plants than in Animals. Plant Physiol. 2005; doi: 10.1104/pp.105.065110.

89. Voichek Y, Hristova G, Mollá-Morales A, Weigel D, Nordborg M. Widespread position-dependent transcriptional regulatory sequences in plants. bioRxiv; 2024; 10.1101/2023.09.15.557872

90. Shore P, Sharrocks AD. The MADS-box family of transcription factors. Eur J Biochem. 1995; doi: 10.1111/j.1432-1033.1995.tb20430.x.

91. Terceros GC, Resentini F, Cucinotta M, Manrique S, Colombo L, Mendes MA. The Importance of Cytokinins during Reproductive Development in *Arabidopsis* and Beyond. Int J Mol Sci. Multidisciplinary Digital Publishing Institute; 2020; doi: 10.3390/ijms21218161.

92. Rose A, Meier I, Wienand U. The tomato I-box binding factor LeMYBI is a member of a novel class of Myb-like proteins. Plant J. 1999; doi: 10.1046/j.1365-313X.1999.00638.x.

